# Machine learning sequence prioritization for cell type-specific enhancer design

**DOI:** 10.1101/2021.04.15.439984

**Authors:** Alyssa J Lawler, Easwaran Ramamurthy, Ashley R Brown, Naomi Shin, Yeonju Kim, Noelle Toong, Irene M Kaplow, Morgan Wirthlin, Xiaoyu Zhang, Grant Fox, Andreas R Pfenning

## Abstract

Recent discoveries of extreme cellular diversity in the brain warrant rapid development of technologies to access specific cell populations, enabling characterization of their roles in behavior and in disease states. Available approaches for engineering targeted technologies for new neuron subtypes are low-yield, involving intensive transgenic strain or virus screening. Here, we introduce SNAIL (Specific Nuclear-Anchored Independent Labeling), a new virus-based strategy for cell labeling and nuclear isolation from heterogeneous tissue. SNAIL works by leveraging machine learning and other computational approaches to identify DNA sequence features that confer cell type-specific gene activation and using them to make a probe that drives an affinity purification-compatible reporter gene. As a proof of concept, we designed and validated two novel SNAIL probes that target parvalbumin-expressing (PV) neurons. Furthermore, we show that nuclear isolation using SNAIL in wild type mice is sufficient to capture characteristic open chromatin features of PV neurons in the cortex, striatum, and external globus pallidus. Expansion of this technology has broad applications in cell type-specific observation, manipulation, and therapeutics across species and disease models.

## Introduction

The biology of the brain is complicated by vast diversity in cell types, subtypes, and cell states. Contemporary advancements in single cell sequencing have identified over a hundred molecularly distinct neuron populations in the mammalian cortex (Hodge et al., 2019; Lake et al., 2016; Saunders et al., 2018; Tasic et al., 2018; Zeisel et al., 2015) including several small subpopulations of Gamma aminobutyric acid (GABA)ergic neurons whose specialized functions are critical for the control of neuronal inhibition (Kepecs and Fishell, 2014; Lim et al., 2018). Understanding neurological function in health and disease from a cell type-specific perspective is critical to the progress of neuroscience.

Such endeavors necessitate cell type-specific technologies for the identification, isolation, and manipulation of discrete cell populations. Transgenic mouse strains targeting major inhibitory neuron subclasses including Parvalbumin-expressing (PV), Somatostatin-expressing (SST), and serotonergic (5-HT) neurons are widely used today and have been instrumental toward our understanding of these cell types (Madisen et al., 2010; Taniguchi et al., 2011). Additional cell type-specific transgenic strains have been created through strategies like enhancer trap (Shima et al., 2016) and EDGE (Nair et al., 2020), which leverage the specificity of *cis* regulatory sequence activity and improve the throughput of transgenic development. Yet even with these innovations, as the number of cell populations of interest rapidly expands, new transgenic strains cannot scale accordingly.

More recently, many developers have turned toward virus-based cell type-specific tools (Dimidschstein et al., 2016; Graybuck et al., 2021; Hrvatin et al., 2019; Mich et al., 2021; Nair et al., 2020; Vormstein-Schneider et al., 2020). Adeno-associated virus (AAV) technologies became particularly attractive with the invention of AAV variants that cross the blood-brain barrier to transduce the central nervous system, AAV-PHP.B and AAV-PHP.eB (Chan et al., 2017; Deverman et al., 2016). In line with certain transgenic engineering, an emerging AAV targeting strategy is to incorporate cell type-specific enhancer elements into the viral genome to promote restricted expression. Enhancer activity can be extremely selective, even more so than the activity of most genes and their associated promoters (Hoffman et al., 2013; Kellis et al., 2014; Roadmap Epigenomics Consortium et al., 2015). Thus, enhancers may be used to confer specificity even for neuron subtypes that cannot be resolved by the expression of a single marker gene (Tasic et al., 2018) or where the marker gene promoter is not specific on its own (Nathanson et al., 2009).

Despite the enthusiasm for enhancer sequences in cell type-specific AAV development, their selection remains nontrivial. ATAC-seq (Buenrostro et al., 2013) has been a popular technique for defining potential cell type-specific enhancer regions because of its high resolution and its compatibility with small cell populations and even single cell technologies (Buenrostro et al., 2015b; Cusanovich et al., 2015). The biggest outstanding barrier to sequence engineering for targeted technologies is the low conversion rate from experimentally suggested cell type-specific open chromatin regions (OCRs) to desired cell type-specific activity in the isolated viral context. Simple enhancer sequence prioritization methods using ATAC-seq signal strength or sequence conservation have been insufficient. Recently, a parallel screening approach involving single nucleus sequencing of barcoded enhancer libraries, PESCA, was proposed to speed up the selection process toward a successful enhancer-driven virus (Hrvatin et al., 2019). Another approach leveraged cell population marker gene proximity for enhancer prioritization (Vormstein-Schneider et al., 2020). We hypothesized that there were additional *in silico* filters that could be applied to reduce the burden of experimental screening in cell type-specific AAV development.

Toward this goal, we sought to leverage the complex combinatorial code linking transcription factor binding site motifs and other DNA sequence features to cell type-specific regulatory activity (Jindal and Farley, 2021). To learn that code, we turned to machine learning models, which have achieved state-of-the-art performance on predicting regulatory activity from DNA sequence (Ghandi et al., 2014; Kelley et al., 2016; Quang and Xie, 2016). Convolutional neural networks (CNNs) (Cun et al., 1989) and support vector machines (SVMs), for example, have been applied to predict enhancer activity from sequence across tissues and cell types (Chen et al., 2018; Kaplow et al., 2020; Kelley, 2020). We reasoned that machine learning classifiers could be applied to identify the most characteristic enhancer sequence patterns within a given cell type, enabling us to prioritize and interpret sequences that are most likely to drive selective expression.

We developed a framework for machine learning-assisted engineering of cell type-specific AAVs, which we refer to as Specific Nuclear Anchored Independent Labeling (SNAIL). Building upon our previously described Cre-activated AAV technology cSNAIL (Lawler et al., 2020), SNAIL probes have the unique advantage of expressing an affinity purification-compatible fluorescent tag (Deal and Henikoff, 2010; Mo et al., 2015). This protein, Sun1GFP, enables nuclei isolation that is particularly advantageous for accessing rare cell populations that would otherwise have low representation in bulk tissue or single nucleus sequencing. Unlike cSNAIL, SNAIL probes are not Cre-dependent, but are instead driven by cell type-specific enhancer sequences selected through machine learning models.

Here, we describe two novel AAV probes for PV neurons. In the mouse cortex, PV SNAIL probes labeled PV neurons with > 70% specificity to Pvalb antibody staining. Isolated populations of tagged cells from the cortex, striatum, and external globus pallidus (GPe) were heavily enriched for known PV open chromatin signatures. In the cortex, PV SNAIL probes were more specific to GABAergic PV interneurons than the common Pvalb-2A-Cre mouse strain. Nucleotide-resolution model interpretation highlighted a collection of 14 transcription factor binding motif families responsible for PV neuron-specific enhancer activation. These results demonstrate concrete utility in sequence-level information for AAV enhancer selection, setting the stage for efficient probe design for a wide range of cell types.

## Results

### Support vector machines discriminate known cell type-specific regulatory sequences

We sought to build machine learning classifiers that could discriminate sequences of differential OCRs between two cell populations. We imposed upfront that training sequences have a minimum fold difference in chromatin accessibility between the cell types to ensure that the model learned cell type-specific features of enhancer activation and not general enhancer features. We chose this strategy because it was most closely aligned with our goal of prioritizing sequences that would activate in one cell type and not others.

To evaluate whether information from differential OCR sequences was sufficient to train accurate classifiers, we first built SVMs comparing select broad classes of cell types in the brain. These were i) a neuron vs. astrocyte classifier and ii) an excitatory neuron vs. inhibitory neuron classifier. The training and validation sequences were based on differential OCRs between cell types, identified from single nucleus (sn)ATAC-seq data from the mouse motor cortex (MOp) (Li et al., 2020) (Supplemental Fig. 1). Both models performed well on held out data, achieving areas under receiver operating characteristic curves (auROCs) of 0.95 and 0.93 (Supplemental Fig. 2).

Next, we verified that these models could recapitulate known cell type-specific activation patterns of commonly used AAV promoter sequences Gfap, CamkII, and Dlx (Supplemental Fig. 2). The Gfap promoter sequence, which empirically has a heavy astrocyte bias *in vivo*, scored highly astrocyte-specific in the neuron vs. astrocyte model, achieving a threshold with less than a 2.1% false positive rate among validation data. In the same neuron vs. astrocyte model, the CamkII promoter and Dlx promoter sequences scored highly neuron-specific. Also consistent with empirical expectations, the excitatory vs. inhibitory neuron model predicted the CamkII sequence to have excitatory neuron preference and the Dlx sequence to have inhibitory neuron preference, while the Gfap promoter scored close to neutral (Supplemental Fig. 2). Therefore, this classification strategy is capable of correctly predicting cell type-specific regulatory sequence activity in the viral context, at least for very distinct cell classes.

### Machine learning models accurately predict PV neuron-specific open chromatin from sequence

Next, we assessed whether the same principles could be applied to more narrowly defined neuron subtypes, using PV neurons as a target. To define potential PV neuron and PV- cell enhancer candidates in the mouse cortex in a data-driven manner, we conducted ATAC-seq on the PV and PV- nuclei populations of Pvalb-2A-Cre mice. The nuclei populations were isolated using previously described Cre-dependent AAV affinity purification technology, cSNAIL (Lawler et al., 2020). cSNAIL probes activate an isolatable nuclear envelope tag in the presence of Cre recombinase protein. Therefore, purified populations from these mice are a direct reflection of cells labeled by the Pvalb-2A-Cre mouse strain, a current standard for PV neuron labeling. These cSNAIL PV and PV- ATAC-seq signatures ultimately defined the training data for models for designing PV SNAIL probes, which are independently activated by PV-specific regulatory elements.

Using merged reproducible ATAC-seq peaks in PV and PV- populations, here called OCRs, we identified significantly differentially accessible OCRs between the two cell populations (DESeq2 padj < 0.01 & |Log2FoldChange| > 1) (Love et al., 2014). To refine these regions for model training, we eliminated promoter-proximal OCRs within 2,000 base pairs (bp) of an annotated transcription start site (TSS). This decision biased training examples toward OCRs of potential enhancer function, which are most relevant for cell type-specific AAV design and may have different sequence composition than gene promoters. This resulted in 14,059 PV OCRs and 4,935 PV- OCRs of interest genome-wide.

We developed two SVMs to distinguish between PV and PV- OCR classes based on nucleotide sequence, one linear model and one nonlinear model. Both SVMs were based on gapped k-mer count vectors, i.e. the number of occurrences of all short subsequences of length k, tolerating some gaps or mismatches, as implemented by LS-GKM (Ghandi et al., 2014; Lee, 2016). The training data were 500 bp sequences underlying PV or PV- OCRs of interest, with a 2.55:1 ratio of positives to negatives. The sequences were centered on ATAC-seq peak summits, where functional transcription factor binding motifs tend to be concentrated (Buenrostro et al., 2013). Taking advantage of this property, we used a center-weighted kernel function for both SVMs, meaning gapped k-mers near the sequence center were weighted more heavily than peripheral gapped k-mers. The two SVMs differed in that one was linear and the other implemented a radial basis function (rbf) kernel, which permits the detection of interactions between gapped k-mers. Both SVMs could predict the correct classification on held out data with high accuracy (Fig. 1b,c), indicating that there were substantial sequence pattern differences between the PV and PV- classes and that the models were able to learn these differences.

**Figure 1:**
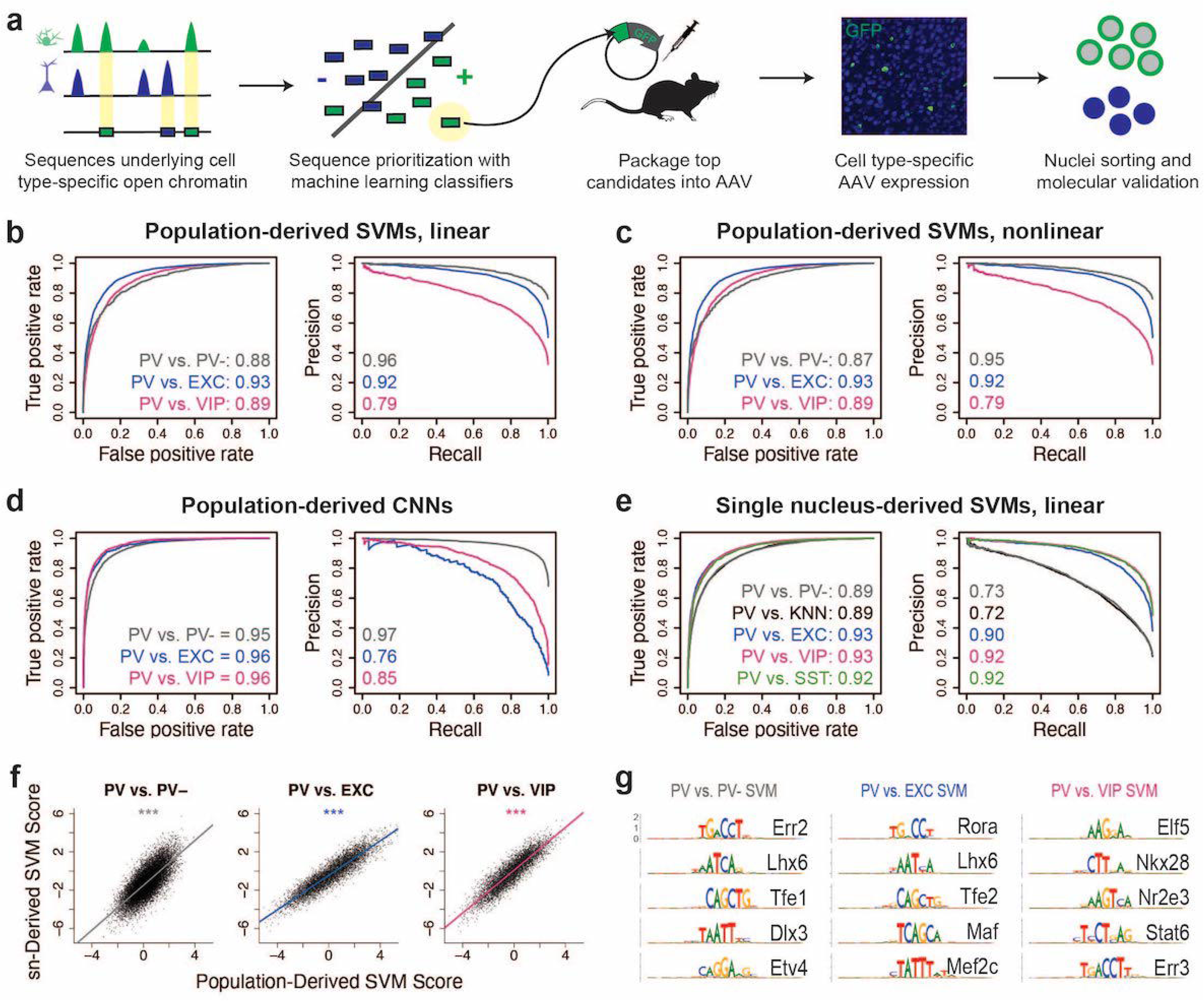
Classification of neuron subtype-specific enhancer activity from sequence. a) Schematic representation of the SNAIL workflow. b-e) Receiver operator characteristic and precision-recall performance metrics for various cell type-specific enhancer sequence model strategies and data modalities. The reported numbers are the areas under the curves for each model. f) Scatter plots for SVM scores reported by equivalent population-derived models and single nucleus-derived models. *** p-value of correlation < 0.001. g) Top five sequence pattern contributors to PV prediction in linear, population-derived SVMs. The best matching known motif is listed (full results in Supplemental Table 12).

Next, because the PV- data contained a high proportion of glial cells, a developmental outgroup to neurons, we considered the possibility that the PV vs. PV- models were learning features of general neuron vs. glia enhancer sequence properties and not necessarily features that were specific to PV neurons. To address this issue, we trained additional population-derived SVMs that directly discriminated between enhancer sequences of PV neurons and other neuron subtypes, using publicly available ATAC-seq data from INTACT-sorted excitatory (EXC) neurons and VIP neurons (Mo et al., 2015). The model training data were defined with the same process described for the PV vs. PV- models. The PV vs. EXC models were trained on 27,879 PV sequence examples and 30,728 EXC sequence examples. The PV vs. VIP models were trained on 15,474 PV sequence examples and 28,683 VIP sequence examples. These models performed well (Fig. 1b,c), indicating that even at the level of neuron subtypes, OCR sequence information is rich enough to reliably distinguish cell type-specific activity.

To survey an additional machine learning strategy, we also built CNN classifiers from the same underlying data, using a different approach (Supplemental Fig. 3). CNNs are best equipped to automatically learn higher-order interactions between sequence features without explicit handcrafting of features. To define the training data for the CNNs, we binned the genome into 200 bp bins and identified bins with differential chromatin accessibility (q < 0.01) between cell types. These sequences were extended bidirectionally to 1,000 bp and used for model training and evaluation. The PV vs. PV- CNN was trained on 55,398 PV sequences and 37,919 PV- sequences, the PV vs. EXC CNN was trained on 3,212 PV sequences and 36,509 EXC sequences, and the PV vs. VIP CNN was trained on 22,416 PV sequences and 96,609 VIP sequences. The CNNs were highly accurate (Fig. 1d), demonstrating an additional approach to discriminate OCR sequence differences between purified neuron populations.

While ATAC-seq from purified cell populations is advantageous for its depth and recovers many examples of differentially accessible reads between neuron subtypes, many neuron populations of interest are not yet isolatable, even through transgenic means. Single nucleus sequencing technologies can be applied to measure neuron subtype-resolution open chromatin without cell sorting by performing several parallel micro-reactions that introduce unique cell barcodes into ATAC-seq sequencing reads. Therefore, we explored whether cell type-specific enhancer sequences derived from mouse motor cortex snATAC-seq (Li et al., 2020) were sufficient to produce neuron subtype-level classifiers. We trained several pairwise linear center-weighted gapped k-mer SVMs to discriminate differential open chromatin sequences from snATAC-seq clusters or groups of clusters. These included analogous models to the population-derived models comparing PV vs. PV-, PV vs. EXC, and PV vs. VIP. In this case, the single nucleus-derived PV vs. PV- model refers to a model trained on differential OCR sequences comparing PV cluster nuclei to all other nuclei with a random sampling probability. The PV vs. k-nearest-neighbor (KNN) model is an additional variation on the PV vs. PV- model where the PV- nuclei sampling for differential OCR analysis was selected for similarity to the PV cluster as implemented in SnapATAC (Fang et al., 2021). We also produced a model comparing PV vs. SST neurons, the most similar subtype to PV. The number of training examples per class of these models ranged from 13,040 to 95,694 and the positive (PV) to negative ratios per model ranged from 1:1.04 to 1:3.74 (further information available in Supplemental Table 2). Single nucleus-derived SVMs were able to classify cell type-specific enhancer sequences with high accuracy (Fig. 1e).

Moreover, models built independently from different data sources identified similar sequence contributions for equivalent tasks. When scoring the population-derived sequences through both the population-derived SVMs and the single nucleus-derived SVMs, individual sequences scored highly similarly in both models (Fig. 1f). These findings highlight the prevalence of reliable cell type-specific enhancer sequence signatures that can be defined by a variety of classifier types and sources of open chromatin measurements. The parameter and performance details of all models can be found in Tables S2 (SVMs) and S3 (CNNs).

### Models learn biological signatures relevant for AAV probe design

We have shown that multiple machine learning strategies are useful for discriminating between regulatory sequences that are differentially active between neuron populations. Next, we asked whether these models could be useful for prioritizing enhancer sequence candidates for cell type-specific enhancer driven technologies. The strength of chromatin accessibility signal at an individual locus may be dynamic and insufficient for cell type-specific enhancer prioritization on its own. Enhancer candidates with highly specific chromatin accessibility and with high specificity scores in the models represent the most characteristic cell type-specific sequence features and may be more effective than other OCRs.

First, we wanted to ensure that the success of the classifiers was rooted in biological sequence signatures related to transcription factor binding motifs. We employed GkmExplain (Shrikumar et al., 2019) and TF-MoDISco (Shrikumar et al., 2018) model interpretation methods to identify sequence patterns with high contributions toward PV neuron-specific OCR predictions, focusing on the population-derived linear SVMs. The models learned sequence patterns that matched known transcription factor binding motifs (Gupta et al., 2007). These included critical developmental transcription factors (TFs) that promote PV interneuron lineage specification *Lhx6*, *Maf*, and *Mef2c (Liodis et al., 2007; Pai et al., 2020; Vogt et al., 2014)* (Fig. 1g). This was encouraging for biological relevance, especially given that the models had no knowledge of known motifs or even the concept of transcription factor binding *a priori*.

To ensure that the neuron subtype-level models were identifying signatures that were relevant for the specific purpose of creating selective PV neuron viruses, we evaluated model predictions on externally validated successful and unsuccessful PV probe enhancer candidates from Vormstein-Schneider et al., 2020, named E1-E34. Importantly, the enhancer sequence from the probe with the lowest PV specificity (E4; 14% specificity) received a negative score from every model, and two probe enhancers with highest cortical PV specificity (E22 & E29; 94% specificity) received high positive scores from every model.

The average score across all models was predictive of probe specificity (Pearson correlation coefficient = 0.42, p = 0.016). Individual enhancer candidates tended to receive similar scores across the SVMs comparing PV to highly abundant cell populations (PV vs. PV-, PV vs. EXC, PV vs. KNN), with Pearson correlations between pairs of models ranging from 0.56 to 0.99 (Supplemental Fig. 4). Many of these models were weakly significant predictors of empirical PV specificity in the AAV context on their own, with the population-derived PV vs. EXC models reaching the highest significance (padj = 0.047) (Supplemental Fig. 5). Some models, such as PV vs. KNN, were better predictors of PV probe specificity than the log fold difference of chromatin accessibility for that cell comparison (Supplemental Fig. 5).

SVMs comparing PV against rarer subtypes (PV vs. VIP, PV vs. SST) were more unique and had less correlation with other models. These models were not significant predictors of probe specificity overall, but many of the highest performing probes had positive scores. Probe specificity was not associated with PhyloP score, which has been considered in cell type-specific enhancer prioritization (Hrvatin et al., 2019), but did show a trend with activity conservation at orthologous regions in the human genome (Supplemental Fig. 5). Importantly, neither method of conservation was as predictive of AAV specificity as the average model score.

This result emphasizes the benefit of enhancer pre-selection with machine learning, which could drastically reduce *in vivo* screening efforts by signaling the best PV enhancer sequences before experimentation. The models predicted which PV enhancer sequence candidates were likely to be cell type-specific drivers and precisely which subsequences were responsible for PV neuron-specific activation. Sequence E29, within the *Inpp5j* locus, was predicted to have PV neuron-specific activity due to a central Mef2 motif site and nearby Err3 motif site, among others (Supplemental Fig. 6). Sequence E22, within the *Tmem132c* locus, was predicted to have PV specificity in part due to Nkx28 and Lhx6 motif sites (Supplemental Fig. 6). Yet, none of these enhancers were our highest predicted PV neuron sequences, so we continued to investigate additional enhancer candidates genome-wide for PV SNAIL probe implementation.

### Two candidate PV SNAIL probes successfully target PV neurons in the mouse cortex

Based on the predictions of all PV enhancer models on our candidates, we prioritized two highly characteristic PV neuron enhancer sequences to test for their ability to drive targeted expression *in vivo* (Fig. 2). We refer to these sequence candidates as SC1 and SC2. Among true PV neuron-specific enhancer sequences that i) were differential OCRs in PV vs. PV-, PV vs. EXC, and PV vs. VIP sorted population data and ii) scored PV positive across all SVM evaluations (1,755 sequences), SC1 was the highest predicted sequence candidate, while SC2 was in the 90th percentile (Fig. 2b, Supplemental Table 4).

**Figure 2:**
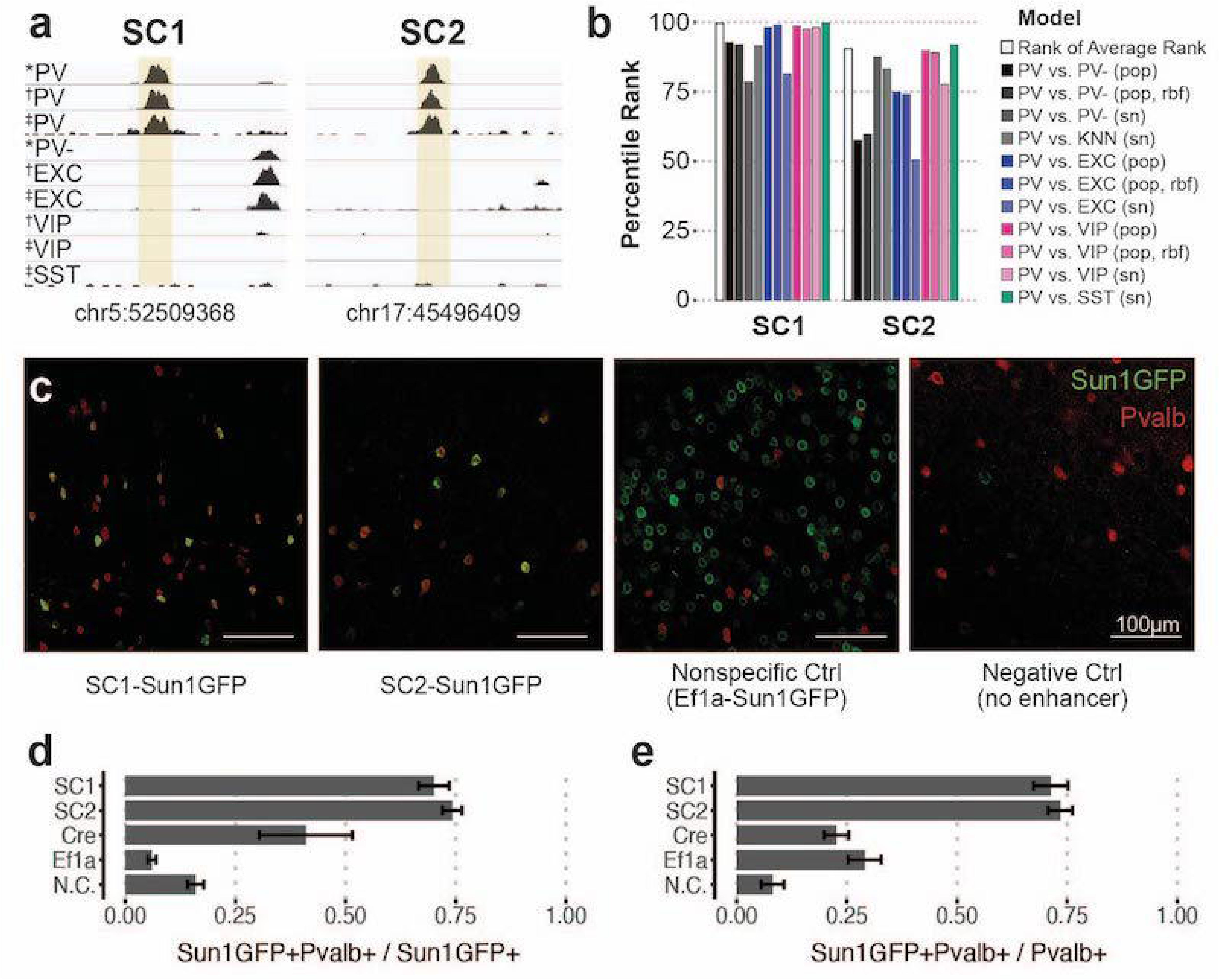
Two sequences candidates selectively activate AAV expression in PV neurons. a) Genome browser visualization of PV specific ATAC-seq signal at sequence candidates SC1 and SC2. * cSNAIL data, † INTACT data from Mo et al., 2015, ‡ snATAC-seq from Li et al., 2020. b) Percentile rank of SVM scores among 1,755 true PV- specific enhancer sequence candidates that scored positively across all models. Linear population-derived models are denoted with “pop”, nonlinear population-derived models are denoted with “pop, rbf”, and linear single nucleus-derived models are denoted with “sn”. c) Example images of AAV Sun1GFP expression against parvalbumin (Pvalb) antibody staining. d,e) Quantification of AAV Sun1GFP or Cre reporter overlap with Pvalb+ cells. Bar heights represent the mean among images and the error of the mean is shown. N cells = 1,322 (SC1), 2,570 (SC2), 1,340 (Cre), 2,013 (Ef1a), and 504 (N.C.). N.C = negative control.

SC1 and SC2 sequences were cloned into separate vectors upstream of the cSNAIL reporter gene, Sun1GFP. To minimize off-target effects, PV SNAIL probes directly rely on transcriptional activation from SC1 or SC2, with no minimal promoter (see methods). We also prepared two control vectors: a negative control that was the identical vector but with no inserted enhancer sequence and a nonspecific control that was the identical vector but with a common Ef1a promoter sequence in place of the candidate sequence. When packaged with AAV-PHP.eB and delivered to the mouse through systemic injection, the SC1-Sun1GFP and SC2-Sun1GFP constructs promoted cortical fluorescence that was restricted to PV neurons, while the Ef1a virus did not (Fig. 2c-e, Supplemental Table 5). Compared with immunohistochemistry-label Pvalb protein, SC1 and SC2-mediated expression of Sun1GFP was restricted to Pvalb+ neurons in ∼70-74% of cases. This was an 11-fold enrichment in precision over the Ef1a promoter and notably, an almost 2-fold enrichment over Cre reporter labeling in Pvalb-2A-Cre mice. We expect these to be conservative estimates of PV targeting due to incomplete antibody capture. On average, Sun1GFP expression from SC1 and SC2 SNAIL probes labeled ∼71-73% of Pvalb+ neurons. The rate is limited by the transduction properties of the AAV-PHP.eB capsid, which only transduces 55-70% of neurons in the cortex (Chan et al., 2017). SC1 and SC2 expression in Pvalb+ neurons represents at least a 9-fold increase over the negative control virus.

### Isolation of PV SNAIL-labeled nuclei captures PV cortical interneurons

Expression of the Sun1GFP gene differentiates SNAIL probes from other cell type-specific AAV technology. The stable nuclear envelope association of this tag enables affinity purification using magnetic beads coated with anti-GFP antibody, which is advantageous for rare population isolation and downstream epigenetic assays. In many contexts, purification of a cell population is more efficient than single nucleus sequencing technologies, especially if the population of interest is in low proportion or the desired downstream applications are not available in single nucleus approaches. Taking advantage of this property, we isolated Sun1GFP-expressing nuclei induced by SC1-Sun1GFP, SC2-Sun1GFP, or Ef1a-Sun1GFP SNAIL virus from the mouse cortex and performed ATAC-seq. Through comparison with known PV neuron ATAC-seq (via cSNAIL in the Pvalb-2A-Cre strain) and PV- or bulk ATAC-seq including cSNAIL PV- cell fractions and Ef1a virus signatures, we determined that both SC1-Sun1GFP and SC2-Sun1GFP cells are highly enriched for PV neurons.

The first principal component, accounting for 84% of the total variance, separated known PV neuron samples from PV- and bulk tissue samples. Likewise, SC1-Sun1GFP and SC2-Sun1GFP samples grouped with the PV samples while Ef1a-Sun1GFP samples grouped with the PV- and bulk sample signatures (Fig. 3a). At the *Pvalb* locus, there were highly reproducible OCR signals between PV cSNAIL, PV snATAC-seq, SC1-Sun1GFP, and SC2-Sun1GFP samples that did not appear in bulk tissue, PV-, or Ef1a-Sun1GFP samples (Fig. 3b).

**Figure 3:**
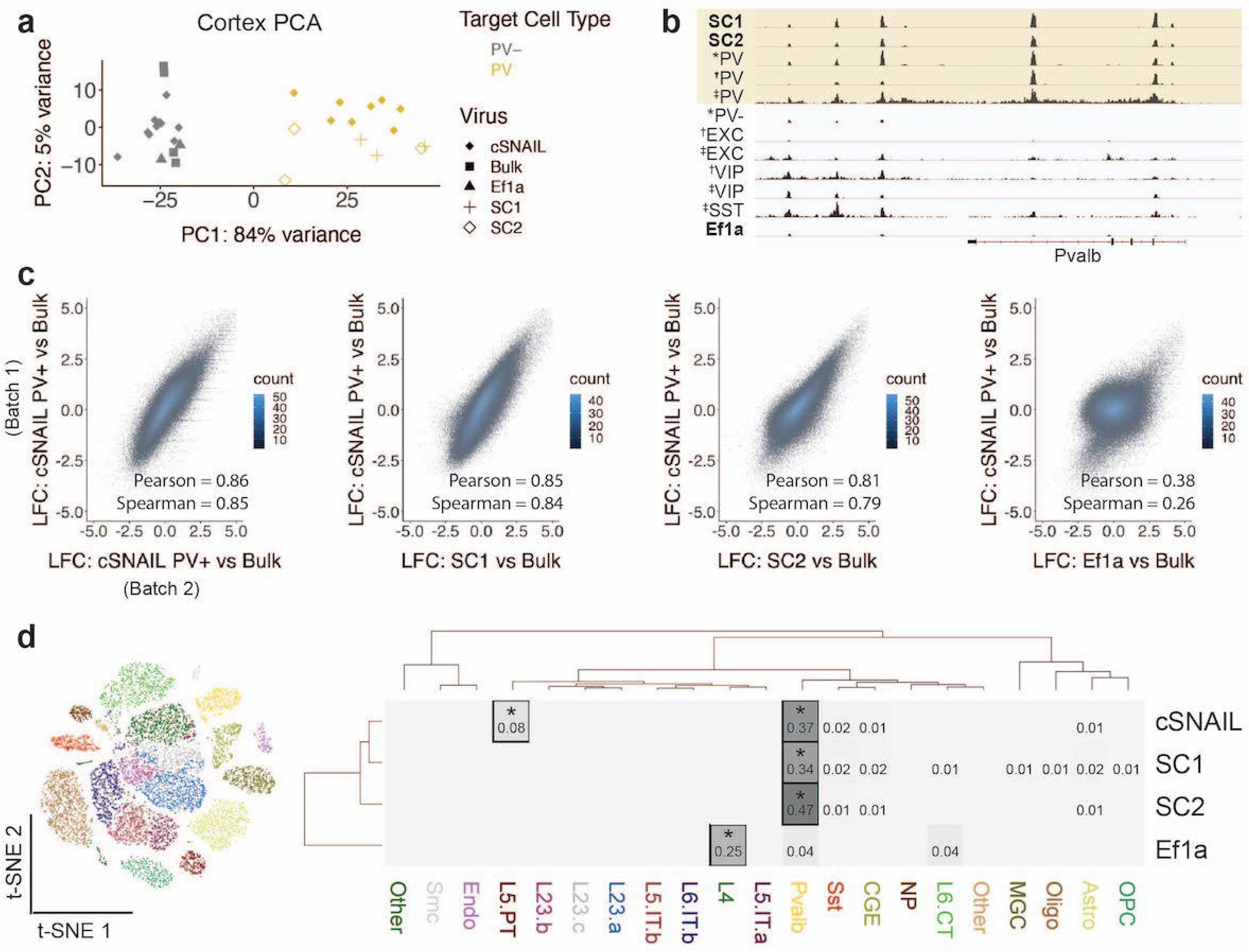
Cortical SC1 and SC2 SNAIL-isolated nuclei recapitulate PV GABAergic interneuron ATAC-seq signatures. a) PCA of ATAC-seq counts across samples. b) Genome browser visualization of ATAC-seq signal at the *Pvalb* gene locus. Tracks represent the pooled sample p-value signal. Each track of similar data type is normalized to the same scale: **SNAIL** data range 0 - 335, *cSNAIL data range 0 - 93, †INTACT data range 0 - 200, ‡snATAC-seq data range 0 - 2. c) Scatter plots of ATAC-seq log2 fold difference relative to bulk tissue ATAC-seq, comparing PV cSNAIL to other AAVs. The density of overlapping points is shown by the plot color. d) snATAC-seq nuclei clusters as visualized by t-SNE. The dendrograms show hierarchical clustering of Euclidean sample distances by Ward’s minimum variance method D2. The heatmap shows the percentage of population OCRs enriched relative to bulk that are also cluster-specific marker OCRs. * Hypergeometric enrichment p < 0.01.

A major goal for PV SNAIL probes was that they may replace transgenic mouse strain technologies in certain contexts. Ideally then, ATAC-seq from Sun1GFP-sorted cells from SNAIL probes in wild type mice should provide similar information as ATAC-seq from Sun1GFP-sorted cells from cSNAIL in Pvalb-2A-Cre transgenic mice. Therefore, we defined PV cSNAIL ATAC-seq log2FoldDifference over bulk cortical tissue ATAC-seq as a gold standard for each OCR. For SC1 and SC2, we computed the correlations between the log2FoldDifference of OCR signal relative to bulk tissue and the log2FoldDifference of OCR signal in PV cSNAIL relative to bulk tissue. To establish an upper limit for correlation, we compared two different batches of cortical PV cSNAIL samples, which had a Pearson correlation of 0.86 and a Spearman correlation of 0.85. As a lower limit, we evaluated the non-specific Ef1a control virus, which had a Pearson correlation of 0.38 and a Spearman correlation of 0.26. Because the AAV-PHP.eB capsid has a neuron bias, these lowly-correlated signatures are likely to be general neuron specifications shared among PV and other neurons. Within this range, SC1 and SC2 had very high correlation with cSNAIL, with SC1 achieving almost equivalent correlation as the two cSNAIL batches (SC1 Pearson = 0.85 and Spearman = 0.84; SC2 Pearson = 0.81 and Spearman = 0.79) (Fig. 3c). The details for differential OCRs in each virus relative to bulk tissue can be found in Supplemental Table 6.

Finally, we compared SC1-Sun1GFP+ and SC2-Sun1GFP+ cell open chromatin signatures to those of snATAC-seq clusters from the mouse motor cortex (Fig. 3d) (Li et al., 2020). We defined cluster-specific OCRs for each snATAC-seq cluster and population-enriched OCRs for SNAIL-isolated cells relative to bulk tissue (see methods) and assessed the overlaps. We found that cSNAIL-isolated PV OCRs, SC1-isolated OCRs, and SC2-isolated OCRs were each significantly enriched for PV cluster-specific markers (34% - 47% overlap, hypergeometric p = 0), while OCRs from Ef1a-isolated cells were not enriched for PV cluster-specific markers (4% overlap, p = 1). Ef1a OCRs instead had the highest enrichment for markers of a layer 4 excitatory neuron cluster (25% overlap, p = 5.3 x 10^-5^). We also note that cSNAIL PV ATAC-seq had an additional 8% overlap with excitatory cluster L5 PT markers (p = 2.5 x 10^-45^), possibly reflective of Pvalb-2A-Cre line labeling in layer 5 Parvalbumin-expressing excitatory neurons (Jinno and Kosaka, 2004; Roccaro-Waldmeyer et al., 2018; Tanahira et al., 2009). These OCRs were absent in SC1- and SC2-isolated cells. In fact, SC1 and SC2 had no enrichment for cluster-specific OCRs of any cluster other than PV (≤ 2% overlap, p > 0.1), including the closely related SST population. This suggests that SC1 and SC2 SNAIL probes actually target a stricter subset of the cells than the Pvalb-2A-Cre mouse strain, likely restricted to PV inhibitory interneurons.

### Chromatin accessibility differences between PV neurons in different brain regions

SC1 and SC2 SNAIL probes were designed based on the sequence properties of cortical PV neurons. Many PV neurons throughout the brain have a common developmental origin in the medial ganglionic eminence (MGE), but there are substantial OCR differences between mature PV neuron populations in different brain regions. From cSNAIL-isolated PV populations in Pvalb-2A-Cre mice (Lawler et al., 2020), we characterized thousands of OCRs with differential accessibility between the cortex, striatum, and GPe (p_adj_ < 0.01, |log2FoldDifference| > 1) (Fig. 4a, Supplemental Table 7). These differences were associated with distinct TF binding motifs (Fig. 4b, Supplemental Table 8). For example, OCRs that were more accessible in cortical PV neurons relative to striatal and GPe PV had highest enrichment for Mef2a motifs, an activity-dependent transcription factor that is important in plasticity and distinguishes subpopulations of PV neurons in the hippocampus (Donato et al., 2015). Mef2c has a similar binding motif and is the second-highest enriched TF motif in cortex-specific PV neuron OCRs. Mef2c is essential for specifying the MGE PV neuron lineage in mouse and human (Mayer et al., 2018) and has been linked to Schizophrenia and other neurodevelopmental disorders (Mitchell et al., 2018). TFs with motifs enriched in PV neuron OCRs that are more open in striatum relative to cortex and GPe included Tgif1, a key homeodomain gene involved in holoprosencephaly (Taniguchi et al., 2012). At 6,654 differential OCRs, GPe-specific PV OCRs were the most unique, and had TF motif enrichments including the Lhx3, Pou5f1, Err3, and Pax3 motifs.

**Figure 4:**
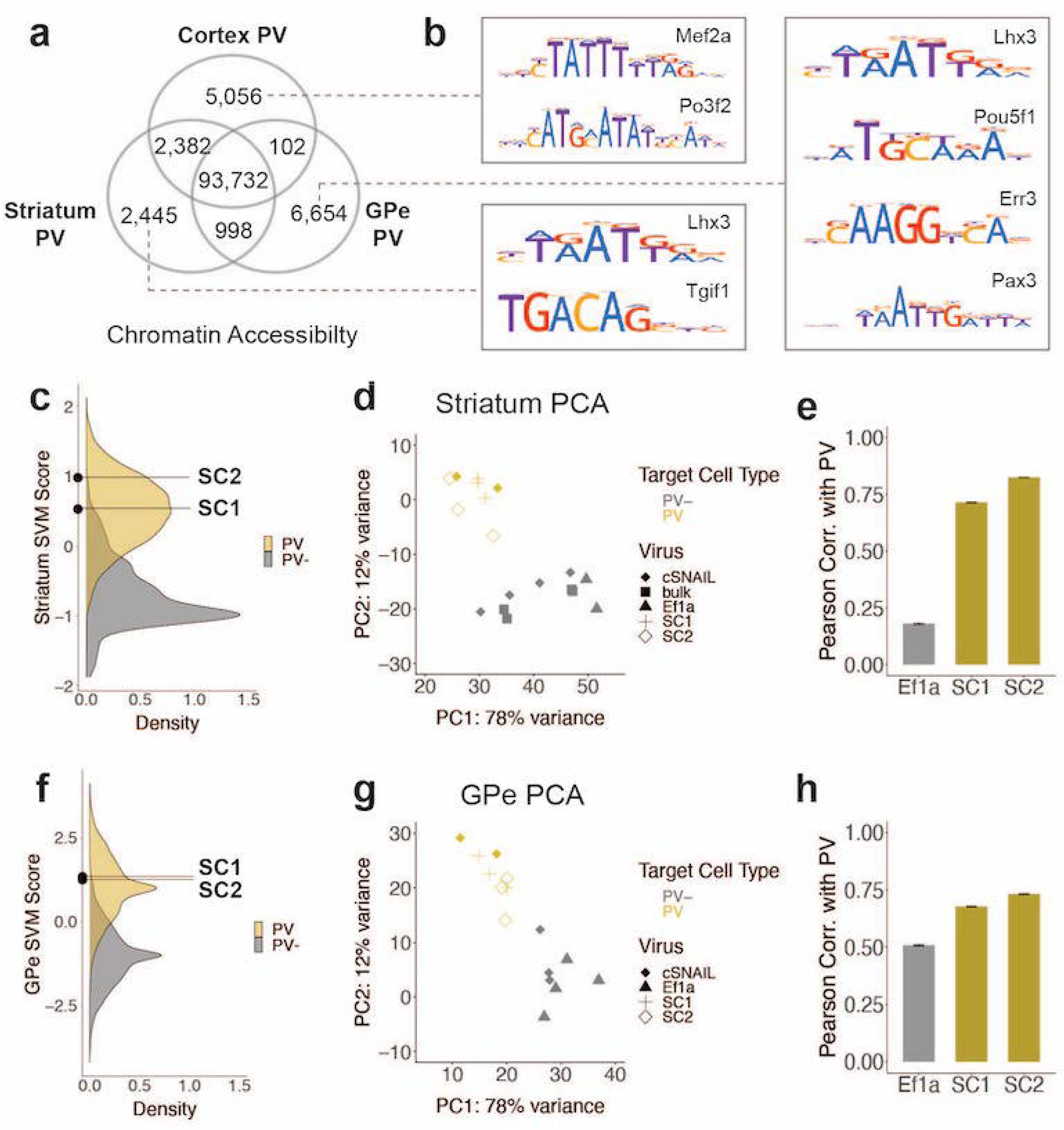
SC1 and SC2 generalize to PV neurons in the striatum and GPe. a) Numbers of differential OCRs between PV neuron populations in three brain regions (DESeq2 padj < 0.01 & |log2FoldDifference| > 1). Brain region-specific OCRs are those that were significantly enriched in that tissue relative to each of the other two tissues. OCRs shared between two brain regions on the venn diagram are those that were significantly enriched in each of those tissues relative to the excluded tissue. The shared center of the venn diagram shows all remaining OCRs that have ambiguous or no tissue preference. b) Examples of enriched motifs in brain region-specific PV open chromatin relative to all PV open chromatin. c,f) Distributions of validation data SVM scores and SC1 and SC2 scores within striatum and GPe PV vs PV- models. d,g) PCA visualization of ATAC-seq counts in each sample. e,h) Pearson correlation coefficients when comparing the log2 fold difference of cSNAIL PV ATAC-seq relative to bulk tissue ATAC-seq and the log2 fold difference of SNAIL ATAC-seq relative to bulk tissue ATAC-seq. Error bars show the 95% confidence intervals.

These molecular differences likely relate to functional differences, for example, the tendency of PV cells in the GPe to project to other brain regions versus the local nature of PV cells in the cortex (Hernández et al., 2015; Saunders et al., 2016). We assessed ontology enrichments in the brain region-specific PV ATAC-seq OCR sets relative to all PV ATAC-seq OCRs using GREAT (McLean et al., 2010) (Supplemental Table 9). The set of PV OCRs enriched in cortical PV neurons included 10 regions associated with the Bdnf gene (Ensembl Genes; FDR Q = 0.0035). Among these was Bdnf promoter IV which is known to be essential for PV neuron synaptic transmission in the prefrontal cortex (Sakata et al., 2009). Other cortex-specific PV enrichments included terms related to sensory perception, especially smell. Striatum-specific PV neuron OCRs were enriched for the adenylate cyclase-inhibiting dopamine receptor signaling pathway (GO:BP; FDR Q = 0.010) and bradykinesia (Mouse Phenotype; FDR Q = 0.046). OCRs preferentially open in GPe PV neurons were enriched for neuropeptide signaling pathways, for example acetylcholine receptor binding (GO:MF; FDR Q = 0.0044) and neuropeptide receptor activity (GO:MF; FDR Q = 1.2 x 10^-5^). This suggests unique epigenetic mechanisms for the regulation of transcription related to receptor signaling in GPe PV neurons, but further work is needed to discern these relationships.

### PV SNAIL probes generalize to subcortical brain regions in the mouse

Given these complexities, we were interested in the extent to which PV enhancer probes chosen from data in one tissue could generalize to other brain regions. Here, we assessed whether SC1 and SC2 SNAIL probes, designed in the cortex, were also selective for PV neurons in the striatum and GPe. First, we used cSNAIL ATAC-seq data from the striatum and GPe to model the regulatory sequence properties of PV neurons vs. PV- cells in these brain regions (Supplemental Fig. 7), and tested whether SC1 and SC2 sequences were predicted to have PV-specific activation (Fig 4c,f). Indeed, SC1 and SC2 were predicted to have PV neuron-specific activity in striatum and GPe PV vs. PV- SVMs. However, there were 1-3,000 sequences with more confident scores toward PV specific activity in each case.

We proceeded to isolate SC1 and SC2-labeled cells from these tissues in wild type mice using Sun1GFP affinity purification and performed ATAC-seq on the tagged populations. We have previously shown high agreement between cSNAIL and Pvalb-2A-Cre labeling in the striatum and GPe (Lawler et al., 2020), so we again used cSNAIL ATAC-seq samples from these regions as true PV neuron signals. By principal component analysis (PCA), we recovered separation between PV samples, including SC1 and SC2-isolated populations, and PV- samples (Fig. 4d,g). We assessed the correlations between log2FoldDifference in SNAIL and cSNAIL samples, each relative to bulk tissue (striatum) or, where there were no bulk samples available, cSNAIL PV- cells (GPe) (Fig. 4e,h, Supplemental Table 10, Supplemental Table 11). Pearson correlation coefficients were similar or slightly lower for SC1 and SC2 in the striatum and GPe than for equivalent comparisons in the cortex, indicating less conservation between cSNAIL and SNAIL probe targets (SC1 cortex = 0.85, striatum = 0.71, GPe = 0.68; SC2 cortex = 0.81, striatum = 0.82, GPe = 0.73). Yet, these were substantially increased over Ef1a correlation with cSNAIL in these tissues, especially for the striatum (Ef1a cortex = 0.38, striatum = 0.18, GPe = 0.51).

By comparing the overlaps of SC1 and SC2-enriched OCRs in striatum and GPe with cortical snATAC-seq cluster-specific OCRs, we still identified the PV cluster as most similar to SC1 and SC2 cells. As expected, all overlaps in striatum-cortex and GPe-cortex comparisons were lower than those from cortex-cortex comparisons, but the magnitudes of SC1 and SC2 overlap with the Pvalb cluster in these brain regions were similar to the magnitudes of cSNAIL PV overlap with the Pvalb cluster in these brain regions (Supplemental Fig. 8). In the striatum, the overlaps with the Pvalb cluster were 8% for SC1, 14% for SC2, and 14% for cSNAIL. In the GPe, the overlaps with the Pvalb cluster were 7% for SC1, 7% for SC2, and 9% for cSNAIL. From these interpretations, SC1 and SC2 SNAIL viruses do generalize to the striatum and GPe, though they may not be as robust as they are within the cortical context.

### Err3 and Mef2 motifs are important for the PV-specific activity of SC1 and SC2 sequences

To interpret the specific sequence patterns within SC1 and SC2 that contribute to their PV neuron-specific activity prediction, we assessed commonly used motifs for each model and identified potential matches within the candidate sequences. For all SVMs, we calculated per-base importance scores and hypothetical importance scores for the set of PV-specific OCRs that were true positives according to all SVMs (score > 0; N = 1,755) (Shrikumar et al., 2019). Then, for each model, we used TF-MoDISco (Shrikumar et al., 2018) to cluster commonly important subsequences called “seqlets” within these PV-specific examples. The resulting clusters represent motifs that were high contributors to a positive score in each model. Among the 11 SVMs comparing PV neuron open-chromatin against PV-cells, EXC neurons, VIP neurons, or SST neurons, we recovered 124 well-supported motifs. Many motifs appeared to be shared across multiple models. Thus, we performed UPGMA clustering on the 124 motifs by sequence similarity using STAMP (Mahony and Benos, 2007) and identified 14 motif clusters (Fig. 5a).

**Figure 5:**
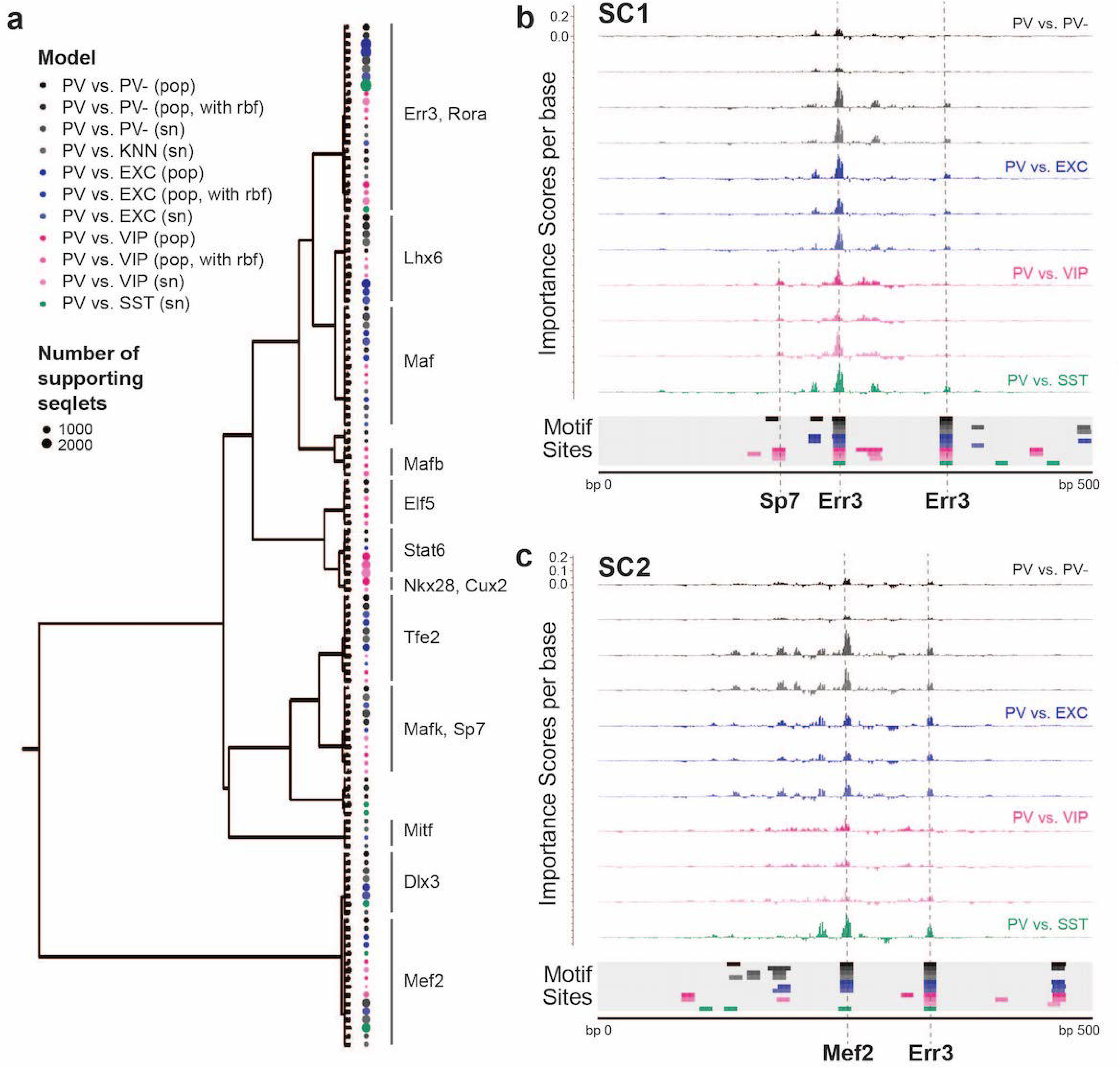
Motif interpretation of PV neuron-specific OCR activity. a) Motifs with high contributions to PV scores in each SVM, clustered by sequence similarity. The bubble color at each node shows the model that motif was discovered in and the size of the bubble shows the number of seqlets supporting that motif. Clusters are labeled by the clade majority best match for known transcription factor binding motifs. The full list of matches can be found in Supplemental Table 12. b,c) Normalized importance of each base in SC1 (b) and SC2 (c) sequences for their PV-specific scores in each SVM. Locations with sequence matches for identified motifs in each SVM (from panel a) are shown at the bottom.

The largest cluster, with 23 motif members, contained representation from all 11 models and had matches to known motifs including the motifs for Err3 and Rora (Supplemental Table 12). Consistent with an important role for Err3 in PV neurons, *Err3* (a.k.a. *Esrrg*) transcript levels were differentially over-expressed in the PV neuron cluster relative the rest of the frontal cortex in snRNA-seq (DropViz subcluster #2-7 Neuron.Gad1Gad2.Pvalb *Esrrg* fold ratio = 8.0, p = 1.14 x 10-198) (Saunders et al., 2018). Esrrg and Rora are key TFs in the Pgc1a transcriptional program, which regulates *Pvalb* expression, mitochondrial function, and transmitter release (Lin et al., 2005; Lucas et al., 2010). Pgc1a signaling is restricted to PV neurons in the brain, and may mediate the unique energy demands of fast-spiking neurons (Lucas et al., 2014; Paul et al., 2017).

The second largest motif cluster contained 16 motifs, also representing all 11 models, and the motifs had best matches to motifs for Mef2a, Mef2c, and Mef2d. In finer subdivisions of this cluster, PV vs. VIP model motifs had best matches to Mef2a, while all other models tended to have best matches for Mef2c and Mef2d. A cluster of Lhx6-like motifs, a transcription factor necessary for MGE interneuron differentiation from interneuron progenitors (Liodis et al., 2007; Vogt et al., 2014), was detected with high support from PV vs. PV- models and PV vs. EXC models, low support from PV vs. VIP models, and not detected between MGE neuron subtypes PV vs. SST. Interestingly, two clusters of motifs were dominated by PV vs. VIP signal, including matches for Stat6, Nkx28, and Cux2 motifs. *Cux2* expression is induced by Lhx6 in the MGE, supporting a role in specification of the MGE interneuron lineage (including PV and SST neurons) from other interneuron lineages (Zhao et al., 2008). Overall, these findings indicate both shared and unique sequence properties dictating PV-specific regulatory sequence activity relative to other cell types.

SC1 and SC2 represent two experimentally validated PV-selective regulatory sequences. To interpret the sequence determinants of their success, we mapped potential motif sites for the 124 TF-MoDISco motifs (Supplemental Table 13) and overlaid these with per-base importance scores for each of the SVMs (Supplemental Table 14). This strategy revealed multiple high importance subsequences with potential transcription factor binding function. SC1 contained two Err3 motifs near the sequence center which were high contributors to the PV-specific model predictions and matched TF-MoDISco motifs for every model (Fig. 5b). An additional subsequence with contributions specific to PV vs. VIP models matched motifs for Sp7. SC2 contained a highly important Mef2 sequence near the center (Fig. 5c). This was a specific match for Mef2c and Mef2d motifs and excluded Mef2a motifs from PV vs. VIP models. Additionally, SC2 contained an Err3 motif with shared importance across all models. Interestingly, the most important features of the SC2 sequence closely resemble those of successful PV probe E29 from Vormstein-Schneider et al., 2020 (Vormstein-Schneider et al., 2020) (Supplemental Fig. 6). The success of SC1 and SC2 are both largely explainable by transcription factor binding motif properties and represent two sequence pattern strategies toward PV-specific activation.

## Discussion

OCR sequence features provide valuable, underutilized information for cell type-specific enhancer design. Here, we showed that sequence alone was sufficient to discriminate between OCR activity in different neuron subtypes. Interpretation of these models revealed rich diversity among the biochemical underpinnings of these classification tasks, reflective of *cis-trans* interactions. The defining sequence properties of cell type-specific OCR activation were robust throughout different data modalities, including ATAC-seq from sorted populations and snATAC-seq, and different classifier types. Machine learning and computational methods, broadly, can facilitate prioritization of AAV enhancer candidates by quantifying sequence properties that are most characteristic and specific to a given cell type.

In SNAIL, our framework for cell type-specific AAV engineering, we incorporate machine learning classifiers as an additional filter for improved enhancer selection. On a set of 33 externally tested PV enhancer-driven AAVs (Vormstein-Schneider et al., 2020), the average PV- specificity score across 11 classifiers was more predictive of PV-specific AAV expression than the log2 fold difference of snATAC-seq signal, sequence conservation, or activity conservation at these loci. With the SNAIL framework, we identified and validated two novel enhancers that drive targeted expression in PV neurons in the mouse cortex. While these do not represent enough trials to establish a new conversion rate from cell type-specific OCRs to cell type-specific AAVs, we were encouraged by the immediate success of the first probes we selected. We believe that incorporation of differential sequence property analyses like those used here will continue to improve the throughput of targeted AAV development in new contexts.

An additional advantage of incorporating classifiers for cell type-specific enhancer selection is increased interpretability of the factors that govern success. The sequence patterns learned by PV models reflected known PV neuron biology. Common motifs contributing to successful PV probe enhancers included Err3, Mef2, and Lhx6, important in the specification and maintenance of the cortical PV interneuron lineage (Liodis et al., 2007; Mayer et al., 2018; Zhao et al., 2008). SC1 and SC2 depend particularly on Mef2 and Err3 motifs for PV specificity.

We found that a combination of multiple direct comparisons between the target cell type and other cell types made for particularly useful screening. Here, we used a tiered approach to ensure specific activity at multiple levels of cellular relationships to PV neurons. At the broadest level, we modeled PV neuron OCR sequences against PV- OCRs, a mixed signature from all other neuron and non-neuron cell types in the mouse cortex. Within neurons, we modeled PV vs. EXC neurons, and then PV relative to more specific subtypes of inhibitory neurons VIP and SST. Successful SC1 and SC2 sequences contained attributes that made them highly PV specific across all of these comparisons.

SC1-Sun1GFP and SC2-Sun1GFP are new AAV technologies for PV neuron labeling and isolation in diverse systems. A unique feature of these viruses is the modified Sun1GFP tag that enables nuclei purification by magnetic beads coated with anti-GFP antibody. This process is advantageous for isolating genomic and epigenomic signals from the population of interest with no dependence on transgenic strains. In comparison to single nucleus sequencing technologies, affinity purification with SNAIL is more efficient for addressing targeted hypotheses about a specific cell type. SNAIL may also be paired with single nucleus sequencing technologies for unprecedented resolution of the substructures within minority cell populations. We took advantage of SNAIL affinity purification to isolate SC1-Sun1GFP and SC2-Sun1GFP nuclei for molecular assessment with ATAC-seq. This represents a novel approach for validating new cell type-specific AAVs. We found that SC1 and SC2 PV SNAIL probes had high molecular agreement with cells tagged in the Pvalb-2A-Cre mouse strain, making them a reasonable alternative to transgenic strain technology. In addition to their success in the intended brain region (cortex), these SC1 and SC2 PV SNAIL viruses also generalized to subcortical regions, the striatum and GPe.

In general, pairing cell type-specific enhancers with AAVs provide much more flexibility and scalability than transgenic technologies. However, there are drawbacks in certain applications. AAVs require time to reach peak expression, usually 2-4 weeks, although some may be robust earlier. This means they are not appropriate for developmental studies in very young animals. Additionally, there are limitations to the transduction efficiency, so AAVs may not be ideal for studies where it is important to label all cells of a certain type. Finally, enhancer activity in AAVs may fluctuate under different ages or in response to different conditions, because enhancers are dynamic actors in the regulation of gene expression. However, machine learning model-based prioritization of characteristic sequences may minimize this risk.

Excitingly, there are many opportunities for extensions of the SNAIL framework that enable cell type-specific interrogation in unprecedented settings. Machine learning model-selected enhancer sequences may be used to drive the expression of a gene for cell type-specific circuit manipulation, as has been achieved with channelrhodopsin and DREADDS (Lee et al., 2010; Vormstein-Schneider et al., 2020). Other important advancements could overexpress a particular ion channel, neurotransmitter receptor, gene variant, or guide RNA for a CRISPR-based gene manipulation strategy. More so than other strategies for cell type-specific AAV design, the SNAIL framework can be tuned for cross-species probe development. In fact, multiple machine learning models have successfully predicted enhancers across mammals, demonstrating high evolutionary conservation in the rules for enhancer sequence activity (Chen et al., 2018; Kaplow et al., 2020; Kelley, 2020; Minnoye et al., 2020). Multispecies models could further improve transferability of probes across species. A new approach that explicitly encourages the model not to learn signatures of species-specific enhancer activity might be especially promising (Cochran et al., 2021). Lastly, while most previous enhancer selection has relied on sorted populations of nuclei from existing transgenic animals, the SNAIL framework provides the opportunity to develop viral tools targeting previously unexplored cell types that are identifiable in snATAC-seq. There is potential to divide subpopulations at multiple levels and design extremely specific technologies. Other applications may exploit changes in enhancer sequence activity in disease and other contexts to target specific cell states. Continued exploration at the intersection of machine learning and enhancer technology development is sure to enhance the impending era of cell type-specific neuroscience and further our general understanding of specific cell types throughout the body.

## Materials and Methods

### Experimental design

The initial cSNAIL experiments to define candidate PV enhancers were performed on primary motor cortex and isocortex samples in triplicate on female mice aged 2-3 months old. All subsequent cSNAIL and SNAIL molecular experiments for the validation of PV SNAIL probes were performed in the cortex, striatum, and GPe with two or three biological replicates. Each of these cohorts included at least one male and one female mouse, all 2-4 months old. Control samples for SNAIL comparisons included cSNAIL PV, cSNAIL PV-, and cells labeled by the Ef1a-Sun1GFP virus. Details for all experiment samples can be found in Supplemental Table 1. Data primary to this publication can be accessed through the NCBI Gene Expression Omnibus (https://www.ncbi.nlm.nih.gov/geo/), accession number GSE171549.

### Nuclei isolation for ATAC-seq

ATAC-seq data were generated using an affinity purification approach with cSNAIL or SNAIL to isolate PV neurons from the mouse isocortex, as described in Lawler et al., 2020. Briefly, mice were overdosed with isoflurane, decapitated, and rapidly dissected. Fresh brain tissue was sectioned coronally on a vibratome for precision, and we dissected brain regions relevant to the specific experiment to be processed as separate samples. All dissections took place in cold, oxygenated artificial cerebrospinal fluid (aCSF). After dissection, we isolated nuclei from the samples by 30 strokes of dounce homogenization with the loose pestle (0.005 in clearance) in lysis buffer as described in Buenrostro et al., 2015 (Buenrostro et al., 2015a). The nuclei were filtered through a 70µm strainer and pelleted with 10 minutes of centrifugation at 2,000 x g at 4 °C. We resuspended the nuclei pellets in wash buffer (0.25 M Sucrose, 25 mM KCl, 5 mM MgCl_2_, 20 mM Tricine with KOH to pH 7.8, and 0.4% IGEPAL) for the affinity purification steps.

### Affinity purification of Sun1GFP+ and Sun1GFP-nuclei

The nuclei suspension was incubated with anti-GFP antibody (Invitrogen, Carlsbad, CA; #G10362) in wash buffer for 30 minutes at 4 °C with end-to-end rotation. After this period, we added Protein G Dynabeads (Thermo Fisher Scientific, Waltham, MA; cat. 10004D) to the reaction and incubated again for 20 minutes. We separated the Sun1GFP+ fraction from the Sun1GFP-fraction on a magnetic bead rack. Sun1GFP-nuclei in the supernatant were centrifuged at 2000 x g for 10 minutes to pellet nuclei, washed one time, and filtered with a 40 µm cell strainer. The Sun1GFP+ nuclei attached to the beads were washed 3-4 times with 800 µL wash buffer by resuspending the sample, letting it settle onto the magnet, and removing the buffer. Where cell yield was not a concern, we also performed a large volume wash with 10 mL wash buffer and filtered through a 20 µm cell strainer. All nuclei preparations were resuspended in water for the ATAC-seq reaction.

### ATAC-seq library construction

For each sample, a small aliquot was stained with DAPI (Thermo Fisher Scientific; cat. 62248) and the concentration of nuclei was determined by counting DAPI+ nuclei with a hemocytometer. Next, we combined 50,000 nuclei, 25 µL Tagment DNA Buffer, and 2.5 µL Tagment DNA Enzyme I (Illumina, San Diego, CA; cat. 20034198) into 50 µL total for the transposition reaction. The reaction incubated at 37 °C for 30 minutes with 300 rpm mixing. Samples containing beads were gently resuspended every 5-10 minutes throughout the incubation to prevent the beads from staying settled at the bottom. Immediately following incubation, the DNA was column purified with the Qiagen MinElute PCR Purification kit (Qiagen, Hilden Germany; cat. 28004). Libraries were amplified to ⅓ saturation with dual-indexed Illumina primers (Preissl et al., 2018). We ensured that samples had the characteristic periodic fragment length distribution of high quality ATAC-seq using TapeStation assessment (Agilent Technologies, Santa Clara, CA). Successful samples were sequenced at low depth on the Illumina Miseq system to determine appropriate library pooling and sequencing depth, then paired-end sequenced for 2 x 150 cycles with the Illumina Novaseq 6000.

### Animal use

All animals for ATAC-seq experiments were either wild type mice (C57BL/6J; Jackson Laboratory, Bar Harbor, ME; Stock No: 000664) for SNAIL experiments or heterozygous Pvalb-2A-Cre mice (B6.Cg-Pvalb^tm1.1(cre)Aibs^/J; Jackson Laboratory Stock No: 012358) (Madisen et al., 2010) on a C57BL/6J background for cSNAIL experiments. Imaging animals were either Pvalb-2A-Cre or double transgenic Pvalb-2A-Cre/Ai14 (Ai14 strain; B6.Cg-Gt(ROSA)26Sor^tm14(CAG-tdTomato)Hze^/J; Jackson Laboratory Stock No: 007914). All mice were 2-4 months old at the time of the tissue experiments. Initial PV cSNAIL data for creating the sorted cell PV vs. PV- model was collected from female mice, but all subsequent validation experiments included representation from both sexes. All animals were housed with a 12 hour light cycle, and experiments were performed 2-3 hours after lights on. Animals for the data primary to this study received no treatments other than the retro-orbital AAV injections. However, previously published cSNAIL data used in analysis included healthy animals that received stereotaxic saline injections to the medial forebrain bundle (Lawler et al., 2020).

### Molecular cloning

To make the non-specific control viral vector pAAV-Ef1a-Sun1GFP, we made modifications to pAAV-Ef1a-Cre with restriction enzyme cloning. pAAV-EF1a-Cre was a gift from Karl Deisseroth (Addgene, Watertown, MA; plasmid #55636; http://n2t.net/addgene:55636; RRID:Addgene_55636). First, we added a multiple cloning site before the Ef1a promoter to create easy promoter swapping for later use. The multiple cloning site insert was synthesized as by Integrated DNA Technologies, Coralville, IA and was inserted between BshTI and MluI sites upstream of the Ef1a promoter. Next, we used BamHI and EcoRI sites to replace the Cre gene with a modified Sun1GFP gene identical to the one in our cSNAIL technologies.

The resulting pAAV-Ef1a-Sun1GFP vector was then further modified to create the other constructs. The PV SNAIL probes were designed to contain one PV-specific enhancer candidate sequence, a synthetic intron for RNA stabilization, the Sun1GFP gene, a WPRE signal, and a polyA signal. From pAAV-Ef1a-Sun1GFP, the Ef1a promoter and intron region was removed and replaced with the sequence for a PV-specific enhancer candidate and the synthetic intron. Inserts for SC1 and SC2 were synthesized by Integrated DNA technologies and cloned into the vector using restriction sites for NdeI and BamHI. To ensure that no expression was being driven from the synthetic intron sequence itself, we similarly cloned a negative control construct containing the synthetic intron, but no enhancer candidate sequence. All transformations during cloning were performed in MegaX DH10B cells (Invitrogen, #C640003) and confirmed with Sanger sequencing.

### AAV production

AAV was produced in AAVpro(R) 293T cells (Takara, Kyoto, Japan; #632273) by co-transfection of the genome pAAV, an AAV helper plasmid, and pUCmini-iCAP-PHP.eB. pUCmini-iCAP-PHP.eB was a gift from Viviana Gradinaru (http://n2t.net/addgene:103005; RRID: Addgene 103005) (Chan et al., 2017). The AAV particles were precipitated with Polyethylene Glycol (PEG 8000, Sigma-Aldrich, St. Louis, MO; cat. P2139-500G) and purified on an iodixanol gradient (OptiPrep, Sigma-Aldrich, cat. D1556-250ML) with ultracentrifugation for 2.5 hours at 350,000 x g at 18°C. We filtered and concentrated the virus in PBS using Amicon Ultra-15 centrifugation filters (Millipore, Burlington, MA; #UFC905024). The viral titer was measured with the AAVpro(R) Titration Kit (Takara, #6233), diluted to a concentration of 8.0 x 10^9^ vector genomes (vg) / µL, and stored single-use aliquots at −80 °C until injection.

### AAV delivery

Animals were anesthetized with 2-3% isoflurane until no pedal withdrawal reflex was observed. Then, we injected 4 x 10^11^ vg total (50 µL) of virus into the retro-orbital cavity and treated the eye with 0.5% Proparacaine Hydrochloride Ophthalmic Solution. The animals were monitored while the virus incubated for 3-4 weeks until endpoint experiments.

### Imaging and analysis

Tissues were fixed with whole body 4% paraformaldehyde (PFA) perfusion and the brains were incubated in 4% PFA for an additional 12-24 hours after dissection. Coronal slices 80 µm thick were made with a vibratome. Free-floating sections were stained for Parvalbumin with Pvalb (Swant, Marley, Switzerland; PV 27) primary antibody with AlexaFluor 405 (Invitrogen, #A-31556) or AlexaFluor 594 (Cell Signaling Technology, Danvers, MA; #8889) secondary antibodies. Images were taken of the motor cortex with laser scanning confocal microscopy. Cells were counted in each channel with Fiji (Schindelin et al., 2012) and assigned as double-labeled or single-labeled manually. Individual images from 1-3 mice were treated as replicates to determine the mean and standard error of the mean for specificity and efficiency quantifications.

### ATAC-seq data processing

Samples were processed from the paired-end fastq files using the ENCODE ATAC-seq pipeline (https://github.com/ENCODE-DCC/atac-seq-pipeline) with the following changes from default behaviors: atac.cap_num_peak = 300000, atac.idr_thresh = 0.1. All samples had high TSS enrichment (>15) and clear periodicity, indicative of good data quality. Optimal IDR peaks were determined for biological replicates of the same cell type, brain region, and sequencing batch (https://github.com/kundajelab/idr) (Li et al., 2011). IDR peaks were then merged to define the combined peak regions (OCRs) for each analysis using bedtools (Quinlan and Hall, 2010). Specifically, we defined sets of OCRs for i) cortex PV and PV- cSNAIL samples, ii) PV, EXC, and VIP INTACT samples (Mo et al., 2015), and iii) cortex, striatum, and GPe bulk samples, PV and PV- cSNAIL samples, SC1-Sun1GFP samples, SC2-Sun1GFP samples, and Ef1a-Sun1GFP samples. We constructed count tables including the relevant samples on each of these OCR backgrounds using Rsubread featureCounts version 1.28.1 (Liao et al., 2019). These three count tables were used to form the basis of i) the sorted population PV vs PV- models, ii) the sorted population PV vs. EXC and PV vs. VIP models, and iii) analysis of SC1 and SC2 SNAIL PV probes in the cortex, striatum, and GPe.

The counts were modeled using the negative binomial distribution in DESeq2 (Love et al., 2014). We assessed the coefficient of cell group, where cell groups were unique tissue, virus, cell type combinations, and we controlled for sex differences where both were present: DESeq2 design ∼ sex + cellGroup. Differential peaks were defined strictly for applications i and ii related to building models (padj < 0.01 and |Log2FoldDifference| > 1) and more loosely for application iii to compare across viruses (padj < 0.05 and |Log2FoldDifference| > 0.5). Related to Fig. 3, only cortical samples from count matrix iii were included in the DESeq2 model, while the Fig. 4 DESeq2 model included samples from all three brain regions.

### snATAC-seq processing

The following samples of snATAC-seq from the mouse MOp were downloaded in Snap file format from http://data.nemoarchive.org/biccn/: CEMBA171206_3C, CEMBA171207_3C, CEMBA171212_4B, CEMBA171213_4B, CEMBA180104_4B, CEMBA180409_2C, CEMBA180410_2C, CEMBA180612_5D, and CEMBA180618_4D(Li et al., 2020). These were processed using SnapATAC version 1.0.0 (Fang et al., 2021). We restricted the analysis to nuclei that passed filtering as defined by the original authors (Li et al., 2020). This removed nuclei that had at fewer than 1000 reads, TSS enrichment <10, or doublet signatures detected by Scrublet (Wolock et al., 2019). Filtered samples contained 6,700-10,983 nuclei each, for a total of 78,525 nuclei. We applied a bin matrix with a bin size of 5,000 and combined the snap objects. Then, we removed bins overlapping with the ENCODE blacklist, mitochondrial regions, and the top 5% of bins that overlapped with invariant features. We reduced dimensionality and selected 18 significant components, then corrected for batch effects using Harmony (Korsunsky et al., 2019). We performed Louvain clustering using runCluster() with the option louvain.lib=”R-igraph”.

Cell types were assigned to clusters by accessibility at promoters and gene bodies of marker genes (Supplemental Fig. 1) and by comparison to the cell annotations from the original authors (Li et al., 2020). Peaks were called for each cluster using MACS2 with the options --nomodel --shift 0 --ext 73 --qval 1e-2 -B --SPMR --call-summits (Zhang et al., 2008). Overlapping peaks across all clusters were merged, resulting in 415,813 OCR regions in total. Differential OCRs were defined using the findDAR() function with test.method = “exactTest” and were required to meet padj < 0.01 (Benjamini-Hochberg corrected) and |log2FoldDifference| > 1. For comparisons to groups of clusters, e.g. PV vs. EXC, separate tests were performed for PV vs. each excitatory cluster, and the intersection of differential OCRs was selected.

### SVM data preparation

SVMs were developed to predict the direction of differential activity from sequences underlying differential OCRs between two cell types or groups of cell types. Because ATAC-seq summit regions are highly enriched for transcription factor binding motifs, we centered on the peak summits within differential ATAC-seq OCRs and extended in both directions for a total fixed sequence length of 500 bp, a convenient length for AAV cloning. Peak summits were defined by MACS2 (Zhang et al., 2008), and only summit regions of peaks called within the cell type of interest were retained. For data from sorted cells, we used optimal IDR peaks across biological replicates of the given cell type. For example, in a PV vs. VIP model comparison, the positive model input examples were 500 bp summit-centered regions of PV IDR peaks that overlapped PV-specific differential open chromatin regions and the negative model input examples were 500 bp summit-centered regions of VIP IDR peaks that overlapped VIP-specific differential open chromatin regions. For snATAC-seq data, we used peaks called within a cluster to define the relevant summit regions. If multiple cell clusters were involved in the comparison, e.g. the excitatory neuron vs. inhibitory neuron model, we used summits found in any peak set from a cluster within that category. In cases where there were multiple summits within a differential open chromatin region, all summits greater than 100 bp apart from each other were retained.

After defining the genomic locations of the summit-centered differential open chromatin regions for model training, we used additional filtering to prepare the data for model training. First, we restricted the models to enhancer regions because they have more specificity than promoters and may be governed by different sequence properties. Therefore, we filtered out regions that were within 2,000 bp of a TSS, using RefSeq annotations downloaded from the UCSC Table browser in July 2020 (Kuhn et al., 2013). Next, we removed super-enhancers because they also may be governed by different sequence features and are not useful for AAV probe design because they are too large. We downloaded mm9 coordinates of mouse cortex super enhancers defined by H3K27ac from the dbSuper database (Khan and Zhang, 2016) and converted these to mm10 coordinates using UCSC liftOver with minmatch = 0.95 (Kuhn et al., 2013). Using bedtools intersect (Quinlan and Hall, 2010), we removed regions with any super enhancer overlap. Finally, we used bedtools getfasta (Quinlan and Hall, 2010) to retrieve the sequences at these genomic coordinates from the mm10 assembly, downloaded from UCSC genome browser in May 2018 (Kuhn et al., 2013), and we removed any sequences that contained uncertain bases (Ns).

### SVM model construction

Sequences were divided into separate partitions by chromosome for model training, validation, and final testing. The training sets included chromosomes 3-7, 10-19, and X, the validation sets included chromosomes 8 and 9, and the test sets included chromosomes 1 and 2. The training data were input into LS-GKM’s gkmtrain and evaluated with gkmpredict (Lee, 2016). Because the input data was summit centered, all models used the center weighted gkm kernel, option -t 4, or the center weighted gkm rbf kernel, option -t 5. The -l, -k, -d, -c, and -w parameters for word length, number of informative columns, number of mismatches to consider, regularization, and class-weighted regularization were tuned to maximize the validation set F1 scores through manual iterations. Other parameters were left on default behavior. auROC and auPRC metrics were calculated and visualized on training, validation, and test sets using the ROCR package in R (http://ipa-tys.github.io/ROCR/). All paper figures reflect final test set performance. The details of all parameter settings and performance metrics of the final models are reported in Supplemental Supplemental Table 2.

### CNN data preparation

We conducted differential accessibility analysis using DESeq2 (Love et al., 2014) to identify regulatory regions that display cell type-specific accessibility in ATAC-seq in PV neurons relative to other background cell types (PV-, VIP, EXC). We used PV and PV- neuron ATAC-seq samples generated in this study as well as PV, VIP, and EXC neuron ATAC-seq samples from Mo et al., 2015. To conduct differential accessibility analysis, we obtained genomic coordinates of all 200 bp bins in the mm10 reference genome, starting from the 200 bp bin at the beginning of each chromosome of including all following contiguous non-overlapping 200 bp bins. We then filtered out any bin that overlaps with an artifact region (Amemiya et al., 2019) or with regions that have unknown nucleotides (obtained from the UCSC twoBitInfo utility using the -nBed option). During this step, regions near the ends of chromosomes were filtered out. Then, using the featureCounts function in the subread package (Liao et al., 2014), we counted the reads mapping to each of the 200 bp bins in the ATAC-seq samples obtained from every included ATAC-seq sample. We then use the DESeq2 R package (Love et al., 2014) to identify bins that were differentially accessible between i) PV and PV-, ii) PV and VIP, and iii) PV and EXC neurons at a Benjamini-Hochberg FDR adjusted p-value cutoff of 0.01. For each of the three comparisons, significant differential bins that displayed PV specificity (log2FoldDifference > 0) were used as positive examples for CNN training and significant differential bins that displayed negative log2FoldDifference (log2FoldDifference < 0) were used as negative examples for CNN training.

### CNN model construction

We trained three separate CNN models that relate sequence to comparative regulatory activity (Kelley et al., 2016; Quang and Xie, 2016; Zhou and Troyanskaya, 2015). For each significant differential 200 bp bin, we obtained the 1000 bp sequence surrounding the center of the bin from the mm10 reference genome and trained the CNN to predict the positive or negative class label. We held out sequence examples underlying all significant differential bins on chromosome 4 as a validation set to evaluate hyperparameter settings and to choose the best performing final model. We also held out sequence examples underlying all significant differential bins on chromosomes 8 and 9 as a test set for final evaluation. Because we had different validation and test sets from those used for the SVM, we did not use any results from the SVM to influence our approach to designing the CNN architecture or any other aspects of CNN training. We implemented our CNN model in Keras 2.2.4 (https://keras.io/) with a theano backend (The Theano Development Team et al., 2016). We created a one-hot encoded representation of the sequence, a 4 x 1000 binary matrix representing positions and occurrences of the 4 nucleotide characters (A,T,G and C) on the sequence, which was propagated through the network. Our CNN architecture consisted of multiple layers of convolution kernels stacked on top of each other (Supplemental Fig. 3). The first such layer consisted of 1000 convolution kernels, each with a kernel width of 8 and height of 4, which scan the input sequence in chunks of 8 nucleotides. We applied rectified linear unit (ReLu) activations on the outputs of these convolution kernels. This initial layer is followed by a variable number of convolution layers with the same number of kernels (100), each of width 8 and height 1. We applied ReLu activations on these convolution outputs as well. These convolution layers are then followed by a set of max pooling operations that selects the maximum value from a set of 13 adjacent units (pooling size = 13). We set the stride for the max pooling operation to 13 units, meaning that it selected the maximum values from contiguous chunks of 13 adjacent outputs from the previous layer. We applied dropout regularization (Srivastava et al., 2014) on the outputs of the max pooling operation to prevent overfitting to the training set. We then flattened the outputs of the max pooling layer into a single vector and passed them to a single output unit with a sigmoid activation function. We used stochastic gradient descent (SGD) to minimize binary cross entropy loss (log loss) between the output of this unit and the positive/negative class label to learn model parameters.

Each model was trained for 100 passes through the training set (or “epochs”). For the PV vs. PV- and the PV vs. VIP tasks, we evaluated model performance and chose the best performing model based on the value of the binary cross entropy loss on the validation set. For the PV vs. EXC task, we chose the final model based on a combination of auROC and auPRC on the validation set. We ignored small differences in validation auROC and auPRC (± 0.02) while selecting the final PV vs. EXC model. Tuning only the number of variable convolution layers (0, 1, or 2), and the dropout probability for the max pooling output (0.2, 0.4, or 0.5), we were able to achieve strong auROCs and auPRCs on the held out validation sets. Therefore, we did not attempt to vary learning rate for SGD (0.01), momentum (0.0), batch size (30), number of training epochs (100), number of filters in the first convolution layer (1000), number of filters in subsequent convolution layers (100), kernel sizes (8), max pooling size (13) and stride (13). A table of hyperparameter settings and associated performance metrics (loss value, auROC, auPRC) on training, validation, and test sets is provided in Supplemental Table 3.

### Broad promoter sequences

The sequences of Gfap, CamkII, and Dlx promoters (Supplemental Fig. 2) were extracted from AAV plasmids with confirmed cell type-specific activity *in vivo*. The Gfap promoter sequence (Gfa2) was from hGFAP-GFP (Addgene plasmid #40592; http://n2t.net/addgene:40592; RRID:Addgene_40592). The CamkII promoter sequence was from pENN.AAV.CamKII0.4.eGFP.WPRE.rBG (Addgene plasmid #105541; http://n2t.net/addgene:105541; RRID:Addgene_105541). The Dlx promoter sequence was from pAAV-mDlx-GFP-Fishell-1 (Addgene plasmid #83900; http://n2t.net/addgene:83900; RRID:Addgene_83900)(Dimidschstein et al., 2016).

### SVM score analysis for external PV AAV screen

33 externally tested PV AAV enhancer sequences (Vormstein-Schneider et al., 2020) were scored through all cortical PV SVMs. To enable comparison between models, scores were normalized to standard deviations from 0 using the standard variation of the validation data set for each model. For each pair of models, the sequence scores were assessed for correlation with cor() function from the R Stats package (https://www.rdocumentation.org/packages/stats/versions/3.6.2) with the Pearson method and visualized using the corrplot package in R (https://github.com/taiyun/corrplot) (Supplemental Fig. 4).

### Alternative prioritization explorations for external PV AAV screen

Common alternative approaches for prioritizing enhancer candidates for cell type-specific AAV design include log2FoldDifference and conservation-based ranking. We show that machine learning models are more predictive of success than these approaches by evaluating on the external PV enhancer AAV screen (Vormstein-Schneider et al., 2020). The log2FoldDifference of ATAC-seq signal in different cell type comparisons was evaluated from snATAC-seq data (Li et al., 2020). We added the exact genomic locations of each test sequence to the genomic peak set for assessment and applied the findDAR() function with test.method = “exactTest” in SnapATAC version 1.0.0 (Fang et al., 2021). The log2FoldDifference was determined for i) the PV cluster relative to all PV- cells using cluster.neg = “random”, ii) the PV cluster relative to closely related cells using cluster.neg = “knn”, iii) the PV cluster relative to the pool of excitatory neuron clusters, iv) the PV cluster relative to the VIP cluster, and v) the PV cluster relative to the SST cluster (Supplemental Fig. 5).

Euarchontoglires PhyloP scores were extracted for all bases within each PV enhancer candidate using the UCSC Table Browser (phyloP60wayEuarchontoGlires track for the Grcm38/mm10 genome, accessed March 2021) (Kuhn et al., 2013). Regions were mapped from mouse (mm10) to human (hg38) using UCSC LiftOver, requiring a minimum ratio of bases that must remap of 0.1. All regions were mappable between species. Finally, we assessed overlapping human PV neuron OCRs from motor cortex snATAC-seq (Bakken et al., 2020) using bedtools intersect (Quinlan and Hall, 2010). Any peak overlap of at least 1 bp was recorded as an overlapping peak.

### Evaluation of SC1 and SC2 ATAC-seq

PCA was performed using plotPCA() on the DESeqDataSet object with variance stabilizing transformation in DESeq2 version 1.26.0 (Love et al., 2014). Using the DESeq2 models described above for cell groups, we extracted OCR statistics for particular cell group comparisons by using the results contrasts. Correlations between log2FoldDifferences for PV cSNAIL vs. bulk tissue and log2FoldDifferences for SNAIL probes vs. bulk tissue were assessed using the R function cor.test() with both “spearman” and “pearson” methods. Genome browser tracks were visualized in the mm10 genome using IGV (Robinson et al., 2011) and track heights were normalized between samples of the same experimental ATAC-seq method (cSNAIL, SNAIL, bulk tissue, or single nucleus). Comparisons to snATAC-seq cluster markers (Fig. 3d, Supplemental Fig. 8) represent the percentage of cSNAIL/SNAIL ATAC-seq OCRs enriched relative to bulk (padj < 0.05 & log2FoldDifference > 0.5) that overlap snATAC-seq cluster markers. snATAC-seq cluster markers were defined as enriched OCRs for that cluster relative to its k-nearest neighbors (padj < 0.01 & log2FoldDifference > 1) that were not enriched OCRs for any other cluster. The significance of the enrichments was assessed using the hypergeometric test with the phyper() function in R, setting lower.tail = FALSE. Enrichments for cluster-specific OCRs were assessed using a background of all snATAC-seq OCRs (N = 415,813) and p-values were corrected for 84 tests with Bonferroni correction.

### Assessment of PV neuron OCRs in different brain regions

PV neuron cSNAIL ATAC-seq samples from cortex, striatum, and GPe tissue of healthy control mice from Lawler et al., 2020 (1 male, 1 female) were assessed for differential open chromatin using DESeq2 as described above. OCRs that were preferentially open in one brain region relative to each of the other brain regions (padj < 0.01 & log2FoldDifference > 1) were evaluated for sequence motif and pathway enrichments. Motif enrichments for tissue-specific PV OCRs were identified using AME version 5.3.3 (Mc Leay and Bailey, 2010) against a background of PV OCRs from all three tissues. Similarly, pathway enrichments using GREAT version 4.0.4 (McLean et al., 2010) were carried out for tissue-specific PV OCRs relative to a background of PV OCRs from all three tissues.

### Model interpretation

We used GkmExplain (Shrikumar et al., 2019) to calculate actual and hypothetical importance scores per base for each of 11 SVMs among 1,755 true positive PV-specific OCR sequences that also scored PV-specific across all SVMs. First, sequences were one-hot encoded. The importance scores were normalized based on the hypothetical importance scores of all possibilities per base, so that a base position decreased in importance if there were other nucleotide possibilities that produced similar scores. We identified sequence motifs with high contributions to PV scores for each SVM separately using TF-MoDISco version 0.4.2.3 (Shrikumar et al., 2018) with options chosen to align with final SVM parameters: sliding_window_size = 7, flank_size = 3, min_seqlets_per_task=3000, trim_to_window_size = 7, initial_flank_to_add = 3, final_flank_to_add = 4, kmer_len = 7, num_gaps = 1, and num_mismatches = 1. The resulting sequence patterns, representing motifs generated from seqlet clusters, were trimmed to the 13 central bases and patterns with support from more than 100 seqlets were used in downstream analysis. The position weight matrices (PWMs) of these patterns were associated with known motifs in the Human and Mouse HOCOMOCO v11 FULL database using Tomtom (Gupta et al., 2007) with the Pearson correlation coefficient motif comparison function (Supplemental Table 12). Motifs from all models were clustered based on PWM similarity using STAMP (Mahony and Benos, 2007); STAMP operations were performed after trimming motif edges with information content less than 0.4, using ungapped Smith-Waterman alignment, the iterative refinement multiple alignment strategy, Pearson correlation coefficient comparison metrics, and UPGMA tree construction. Finally, individual instances of motif sites were mapped in SC1 and SC2 sequences using FIMO with default parameters (Grant et al., 2011).

## Acknowledgements

This material is based upon work supported by NIH Grant DP1DA046585 and the National Science Foundation Graduate Research Fellowship Grants DGE1252522 and DGE1745016.

