## Supplemental Figure and Table Legends for "Machine learning sequence prioritization for cell type-specific enhancer design"

Supplemental Figure 1: snATAC-seq cluster assignments. a) Cluster annotations in tSNE space. b) Gene body accessibility of population marker genes. Data reprocessed from Li et al., 2020.

Supplemental Figure 2: Validation of modeling strategy using known promoters targeting broad cell classes. a) Performance metrics for SVMs trained to distinguish differential OCRs between Neurons vs. Astrocytes or Inhibitory neurons vs. Excitatory neurons. b) Model scores for Gfap, CamkII, and Dlx cell type-specific promoter sequences.

Supplemental Figure3: CNN strategy overview.

Supplemental Figure 4: Correlations between SVM scores across models. Scores were evaluated on 33 enhancer sequences from Vormstein-Schneider et al., 2020

Supplemental Figure 5: Model predictions on experimentally validated PV AAV enhancer candidates. a) Model scores for enhancer candidates, normalized as standard deviations from 0 based on model validation sequence scores. The right panel indicates the significance of the correlation between specificity *in vivo* and SVM scores for each model. b) log2 Fold Differences in enhancer accessibility between the PV cluster and a KNN background set in snATAC-seq data (Li et al., 2020). The right panel indicates the significance of the correlation between specificity *in vivo* in the AAV context and log2FoldChange in snATAC-seq. For a) and b), p-values (except for the average hypotheses) were corrected for multiple hypothesis testing with the Benjamini Hochberg method. c) Distributions of phyloP conservation scores across EuarchontoGlires for nucleotides within enhancer regions. The median score per enhancer is shown by the red point. The bottom of the panel shows presence (+) or absence (-) of an overlapping human OCR from human Snare-seq data (Bakken et al., 2020).

Supplemental Figure 6: Interpretation of top external PV AAV enhancer sequences. Normalized importance scores per base of PV enhancer candidates E29 and E22 across linear, population-derived SVMs. The locations of TF-MoDISco motif sites are shown at the bottom of each panel.

Supplemental Figure 7: Subcortical PV vs. PV- SVMs. ROC and PRC performance metrics on held out test sequences are shown.

Supplemental Figure 8: Comparison of subcortical SC1 and SC2-labeled populations with cortical snATAC-seq cluster markers.

Supplemental Table 1: Sample metadata information.

Supplemental Table 2: SVM parameter tuning and performance evaluations.

Supplemental Table 3: CNN parameter tuning and performance evaluations.

Supplemental Table 4: Model predictions across 1,755 experimental PV-enriched OCRs with positive PV model scores.

Supplemental Table 5: Image quantification for SNAIL viruses or the Pvalb-2A-Cre mouse strain reporter with Pvalb immunohistochemistry.

Supplemental Table 6: Differential OCR statistics for PV SNAIL-isolated cortical ATAC-seq relative to bulk tissue cortical ATAC-seq.

Supplemental Table 7: Differential OCR statistics for tissue-specific PV neuron OCRs in cortex, striatum, and GPe.

Supplemental Table 8: Motif enrichments among tissue-specific PV neuron OCR sequences.

Supplemental Table 9: Pathway enrichments among tissue-specific PV neuron OCR sequences.

Supplemental Table 10: Differential OCR statistics for PV SNAIL-isolated striatal ATAC-seq relative to bulk tissue striatal ATAC-seq.

Supplemental Table 11: Differential OCR statistics for PV SNAIL-isolated GPe ATAC-seq relative to PV- GPe ATAC-seq.

Supplemental Table 12: Association of TF-MoDISco motifs to known transcription factor binding motifs.

Supplemental Table 13: Identification of motif sites within SC1 and SC2.

Supplemental Table 14: Normalized per-base importance scores for SC1 and SC2 sequences.
